# Multimodal analyses reveal genes driving electrophysiological maturation of neurons in the primate prefrontal cortex

**DOI:** 10.1101/2023.06.02.543460

**Authors:** Yu Gao, Qiping Dong, Kalpana Hanthanan Arachchilage, Ryan D. Risgaard, Moosa Syed, Jie Sheng, Danielle K. Schmidt, Ting Jin, Shuang Liu, Sara Knaack, Dan Doherty, Ian Glass, Jon E. Levine, Daifeng Wang, Qiang Chang, Xinyu Zhao, André M. M. Sousa

## Abstract

The prefrontal cortex (PFC) is critical for myriad high-cognitive functions and is associated with several neuropsychiatric disorders. Here, using Patch-seq and single-nucleus multiomic analyses, we identified genes and regulatory networks governing the maturation of distinct neuronal populations in the PFC of rhesus macaque. We discovered that specific electrophysiological properties exhibited distinct maturational kinetics and identified key genes underlying these properties. We unveiled that RAPGEF4 is important for the maturation of resting membrane potential and inward sodium current in both macaque and human. We demonstrated that knockdown of *CHD8*, a high-confidence autism risk gene, in human and macaque organotypic slices led to impaired maturation, via downregulation of key genes, including *RAPGEF4*. Restoring the expression of *RAPGEF4* rescued the proper electrophysiological maturation of CHD8-deficient neurons. Our study revealed regulators of neuronal maturation during a critical period of PFC development in primates and implicated such regulators in molecular processes underlying autism.

## INTRODUCTION

The dorsolateral prefrontal cortex (dlPFC), a neocortical area highly derived in primates^1–3^, is required for higher-order brain functions, including executive functions such as working memory, planning, and decision-making^3–5^. These functions are significantly impaired in brain disorders, including autism spectrum disorders (ASD), schizophrenia, obsessive-compulsive disorder, and neurodegenerative disorders^6–8^. Much of our knowledge on PFC development is largely extrapolated from rodent studies. Although rodents possess a PFC that is homologous to a small portion of the primate PFC, they lack the granular dlPFC^1^. Several comparative studies have identified gene expression signatures that are unique to the PFC of humans and non-human primates (NHP)^3, 9–12^. Therefore, it is essential to understand the mechanisms regulating dlPFC development in primates.

The cells that compose the primate dlPFC, especially excitatory and inhibitory neurons, undergo extensive and dynamic maturation throughout midfetal and late-fetal development, during which critical neurodevelopmental events, such as circuit assembly and electrophysiological maturation of neurons occur^13^. Midfetal and late-fetal development are a convergent period of expression for many ASD genes^14, 15^, yet the functions of these ASD genes during this period of development remains unclear. Furthermore, although we have a robust knowledge of the molecular and cellular mechanisms that govern cell fate specification during cortical development^16–22^, and of the transcriptomic and electrophysiological properties of mature neurons in the adult neocortex^23–27^, our understanding of the gene networks that regulate the early stages of neuronal maturation, especially electrophysiological maturation, remains elusive. Therefore, a multimodal investigation of primate dlPFC neurons that combines electrophysiological and functional genomic analyses during critical periods for neuronal maturation is essential to understand the mechanisms driving neuronal maturation.

Here, to uncover the molecular mechanisms that regulate the maturation of dlPFC neurons, we first generated single-nucleus multiomic (snMultiomic) data and performed integrated analyses of the dlPFC from rhesus macaques (*Macaca mulatta)*, ranging from midfetal to late-fetal periods. We then performed Patch-seq on acute dlPFC slices from 16 macaques, including the same specimens profiled for snMultiome. This approach allowed us to integrate our electrophysiological analyses with gene expression profiles to identify the genes that are important for the maturation of specific electrophysiological features. Furthermore, we have evaluated the function of select genes – *RAPGEF4* and *CHD8 –* and demonstrated that a loss of function of these genes affects the morphological and electrophysiological maturation of cortical neurons in organotypic slices of macaque and human. Finally, Patch-seq analysis of brain slices with *CHD8* knockdown identified RAPGEF4 as a key mediator for CHD8 regulation of electrophysiological maturation of human cortical neurons.

## RESULTS

### Cell type-specific transcriptomic changes underlying the maturation of rhesus macaque dlPFC cells during midfetal and late-fetal development

We performed snMultiomic analyses on 9 rhesus macaque dlPFC samples across eight prenatal ages: postconceptional day (PCD) 85, PCD95, PCD100, PCD105, PCD110, PCD125, PCD145, and two PCD155 (**Figures 1A-1B; Table S1**). After implementing stringent quality control measures, we retained 76,855 nuclei for further analyses (**Figures S1; Tables S1-S2**). After removing batch effects, we used unsupervised clustering to group nuclei into transcriptomically defined groups (**Figures S1B-S1C**). Using this strategy, together with cell type-specific marker genes^16, 23, 28^, we defined 9 major cell classes that included glutamatergic excitatory neurons (ExN), GABAergic inhibitory neurons (InN), glial progenitor cells (GPC), astrocytes, oligodendrocyte precursor cells (OPC), oligodendrocytes, microglia, vascular leptomeningeal cells (VLMC), and endothelial cells (**Figure S2A**). We re-clustered the ExN and InN nuclei to identify their subclasses based on the layer (L) and projection identity (intratelencephalic [IT], extratelencephalic [ET], near-projecting [NP], corticothalamic [CT], and L6B) for ExN, and the developmental origin (medial ganglionic eminence [MGE], caudal ganglionic eminence [CGE], or dorsal lateral ganglionic eminence [dLGE]) for InN (**Figures 1C and S2B-S2C**). Interestingly, one small population of IT neurons exhibited molecular features of both ExN L2-3 IT and ExN L3-5 IT, indicating that this small population of neurons is yet to be fully specified; we classified this population as ExN L2-3 IT/L3-5 IT. At the lowest hierarchical level of cell type classification, we transcriptomically defined 85 cell subtypes (**Figure S2E; Table S3**). By integrating our dataset with two published snRNA-seq datasets that analyzed early fetal^28^ and adult^23^ macaque dlPFC, we observed that our data bridges the embryonic/early fetal cell types with the adult cell types, facilitating the tracing of the developmental origins of distinct adult cell subtypes (**Figures 1E-1G**).

**Figure 1.**
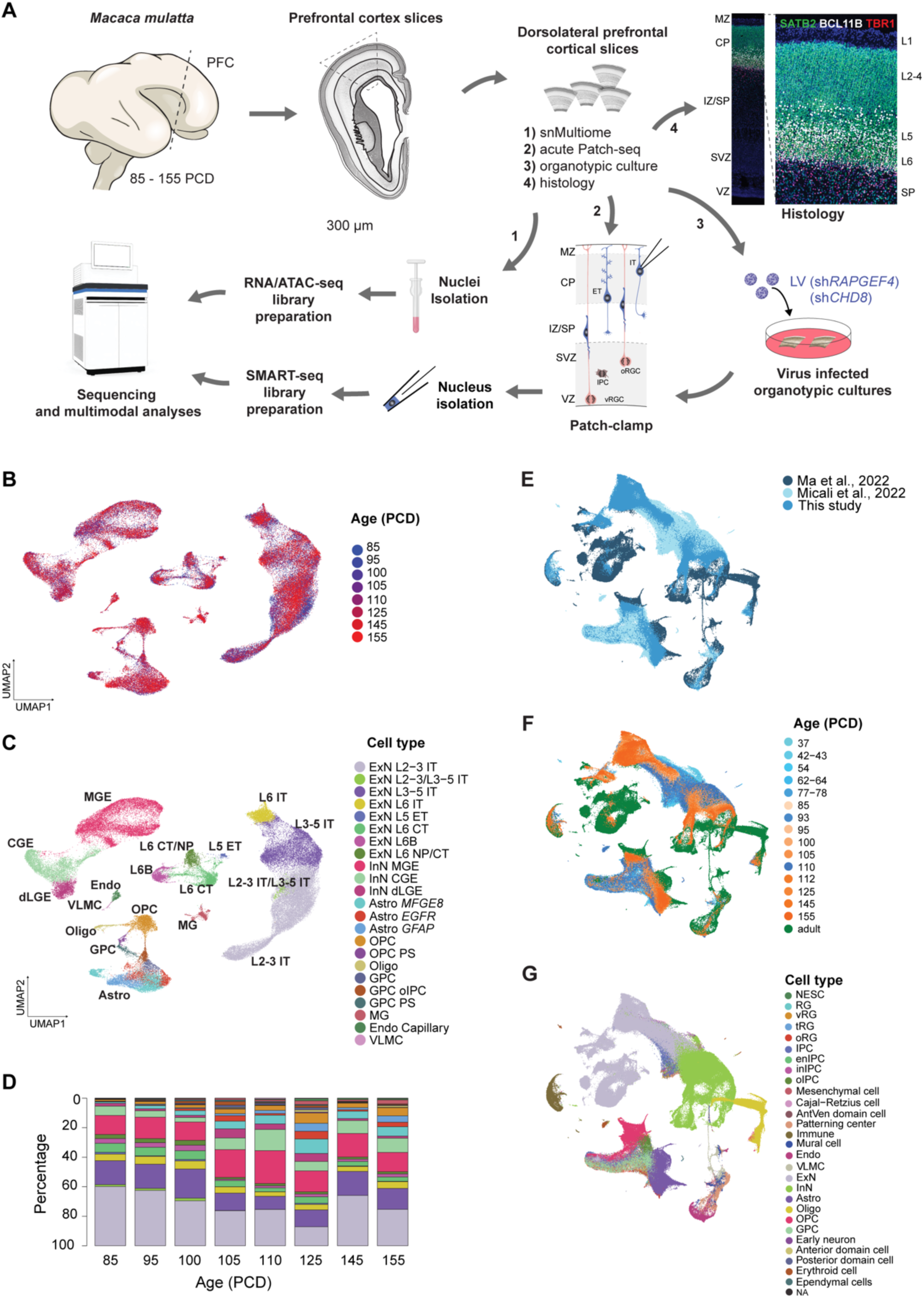
Single-nucleus transcriptomic atlas of rhesus macaque dlPFC cells across midfetal and late-fetal development. (A) The frontal cortex of Rhesus macaque (*Macaca mulatta*) brains, aged from postconceptional day (PCD) 85 to 155, were dissected and sectioned at 300 μm. Acute dorsolateral prefrontal cortices were analyzed using Patch-seq and single-nucleus multiome (snRNA-seq and snATAC-seq). Tissue from the same specimens, as well as human midfetal tissue, was also used to culture organotypic slices for functional analyses of select genes (*RAPGEF4* and *CHD8*). Validation of key findings was also performed by multiplexed immunostaining and single-molecule fluorescent in situ hybridization on dlPFC sections from the contralateral hemisphere of the same specimens (**Table S1**). Finally, we have performed an integrative analysis of the generated multimodal dataset. (B-C) UMAP visualization of all the cell classes and subclasses. Cells are colored according to age in postconceptional days (PCD), from PCD85-PCD155 (B) cell subclass (C). (D) Bar plot showing the cell composition of the dlPFC throughout midfetal and late-fetal development. (E-G) Integrated UMAP visualization depicting cells obtained from both the current study and published datasets^22, 27^. Cells are color-coded based on their respective dataset (current study in medium blue) (E), age of donor specimen (embryonic/early fetal in blue, current study in orange, adult in dark green) (F), and cell type (G). ExN, excitatory neuron; InN, inhibitory neurons; Astro, astrocytes; Oligo, oligodendrocytes; IPC, intermediate precursor cells; enIPC, excitatory neuron IPC; inIPC inhibitory neuron IPC; oIPC, oligodendrocyte IPC; OPC, oligodendrocyte precursor cells; GPC, glial precursor cells; Endo, endothelial cells; RG, radial glia cells; oRG, outer radial glia cells; tRG, truncated radial glia cells; vRG, ventricular radial glia cells; NESC, neuroepithelial stem cells; VLMC, vascular leptomeningeal cells; AntVen domain cell, antero-ventral domain cells; NA, unknown cell type.

As expected, we observed an increasing number of glial cells, including astrocytes, OPCs, and oligodendrocytes throughout late-fetal development (**Figure 1D**), a period characterized by minimal cortical neurogenesis and robust gliogenesis^29^. Within the excitatory neurons, we observed that IT neurons were more abundant than any other subclass, as previously described in the adult dlPFC^23^. The selective expansion of IT neurons in the primate lineage, especially of upper-layer (L2-3) neurons^23^ and their functional relevance in the cortico-cortical circuits that govern some of the unique cognitive and behavioral abilities of primates^30^, led us to focus our analysis on the maturational profiles of these IT neuronal populations: ExN L2-3 IT and ExN L3-5 IT (**Figures 2A-2D**). We performed differential gene expression analysis along the developmental trajectory (i.e., pseudotime) to identify genes that display dynamic, highly variable changes during neuronal maturation. Among the top 100 genes (**Table S4**), we observed genes that exhibit cell type-specific correlation with maturation (**Figures 2C-2D**), including *MEIS2*, a transcriptional regulator that is induced by retinoic acid and is important for the arealization of the prefrontal cortex^10^, in L2-3 IT, and *FOXP2*, a gene encoding a transcription factor that has recently been shown to have primate-specific expression in L3-5 IT excitatory neurons^23^. Disruptions in these genes have been linked to neuropsychiatric disorders^31, 32^.

**Figure 2.**
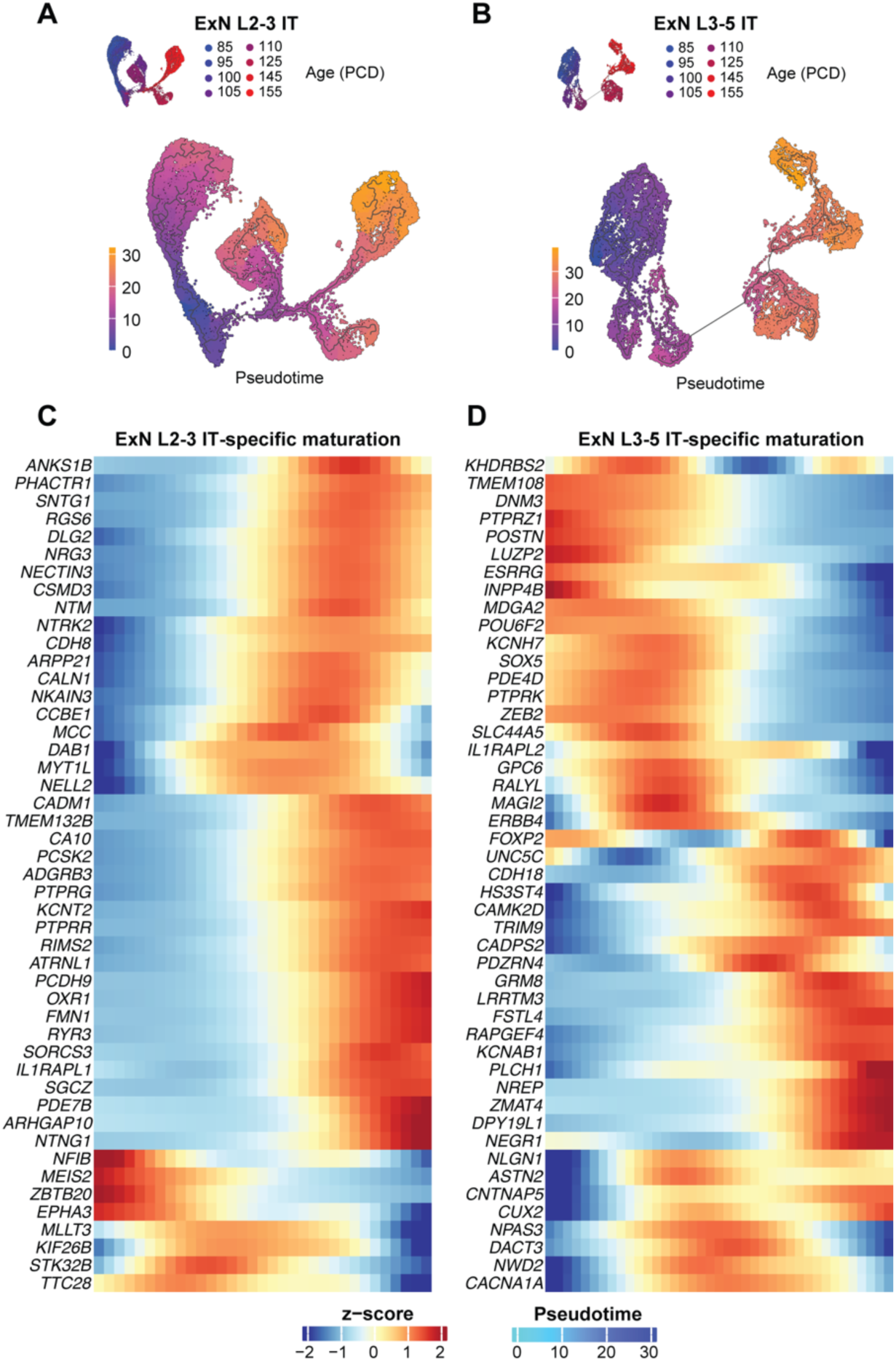
Maturational trajectories of L2-3 and L3-5 intratelencephalic-projecting neurons. (A-B) UMAP visualization of the maturation trajectory of ExN L2-3 IT (A), and ExN L3-5 IT (B) cells; the top panel is colored according to postconceptional days, whereas the bottom panel is colored according to inferred pseudotime. (C-D) Gene expression variation with pseudotime of genes specific to ExN L2-3 IT (C), ExN L3-5 IT (D) (from the top 100 highly variable genes along the developmental trajectory).

Together, our analyses uncovered the transcriptional landscape of primate midfetal and late-fetal PFC development, highlighting differences between distinct excitatory neuronal populations throughout maturation.

### Chromatin accessibility signatures and gene regulatory networks of macaque midfetal and late-fetal dlPFC cells

The simultaneous profiling of gene expression and chromatin accessibility in the same cells allows linking putative regulatory elements to genes, thus providing insights into cell-type gene regulatory mechanisms in the developing primate brain. After filtering out low-quality nuclei and implementing other stringent quality control steps, snATAC-seq data from 64,941 nuclei were retained for downstream analyses (**Figure S1D**). We observed that the clustering of cells based on accessible chromatin was similar to the clustering utilizing gene expression data (**Figures S1E-S1H**), showing the congruence between the two datasets (**Figure 3A**). However, the snATAC-seq data revealed more pronounced differences in the maturation states of cells within each cell subclass, especially for IT excitatory neurons (**Figure S1E**). To gain a better understanding of the association of regulatory elements and their target genes during neuronal maturation, we estimated peak-to-gene links, which identify co-activity between genes and their nearby accessible chromatin regions, for the excitatory and inhibitory neurons (**Figure S3A**), thus providing a list of potential regulatory elements for each gene (**Table S5**). We found that peak-gene combinations were hierarchically divided into three distinct categories: a cluster specific to excitatory neurons during early midfetal development, a cluster specific to excitatory neurons during late-fetal development, and a cluster specific to inhibitory neurons during late-fetal development. Functional enrichment analysis of each group revealed that the clusters specific to late-fetal development were enriched for terms related to calcium signaling and dendritic development in excitatory neurons, and synaptic processes in inhibitory neurons, whereas the early-midfetal development cluster was enriched for transcriptional regulation terms (**Figures S3C-S3E; Table S5**).

**Figure 3.**
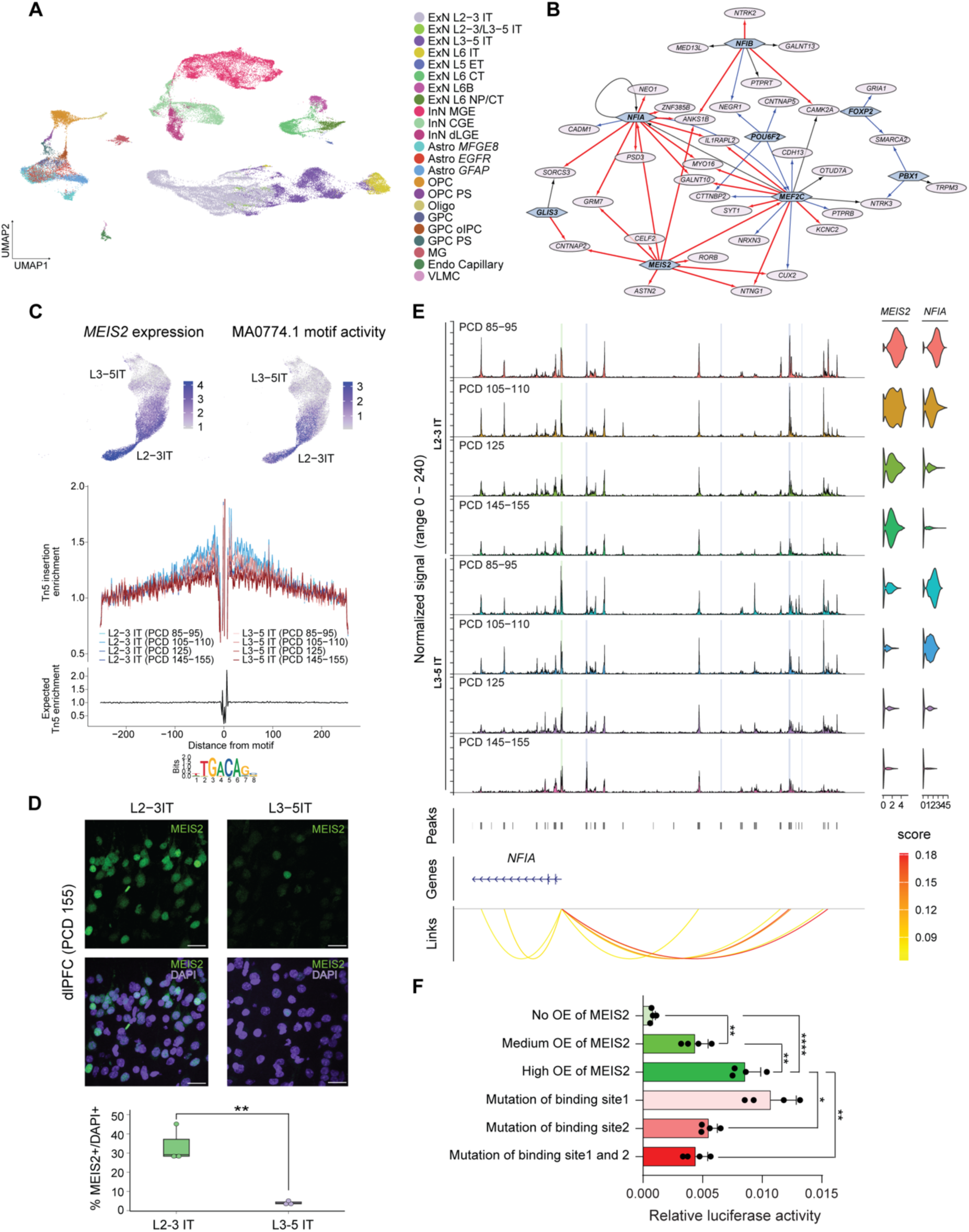
Single-nucleus multiome gene expression and chromatin accessibility of rhesus macaque dlPFC cells across mid- to late-fetal development. (A) Joint (snRNA-seq and snATAC-seq) UMAP visualization of cell subclasses. (B) Differential gene regulatory networks inferred for ExN L2-3 IT and ExN L3-5 IT cell subclasses. Hexagonal nodes indicate transcription factors, and the elliptical nodes indicate the target genes. The edge colors represent the cell subclass specificity of transcription factor-target gene relationships (black, conserved in both cell subclasses; red, ExN L2-3 IT specific; blue, ExN L3-5 IT specific). (C) UMAP visualization of *MEIS2* expression and motif activity scores for MA0774.1 (MEIS2 transcription factor motif) in L2-3 IT and L3-5 IT cells (top panels); MEIS2 transcription factor footprints for ExN L2-3 IT and ExN L3-5 IT cells across four developmental periods (bottom panel). (D) Immunofluorescent detection of MEIS2 in the macaque dlPFC at PCD155, in L2-3 IT and L3-5 IT cells shows that a significantly higher number of cells express MEIS2 in L2-3 IT than in L3-5 IT cells (L3-5 was identified using in situ hybridization against *RORB*, as shown in Figure S3E). Scale bar, 20 µm. (**p < 0.01, *t*-test). (E) Peak-to-gene link plot for *NFIA* together with the gene expression of *MEIS2* and *NFIA* in ExN L2-3 IT and ExN L3-5 IT across four developmental periods. The links indicate the co-activity links between *NFIA* gene expression and chromatin accessibility peaks. The highlighted regions within the chromatin accessibility peaks show the MEIS2 transcription factor motif (MA0774.1) binding sites. (F) Dual-Luciferase reporter assay of *NFIA* genomic sequences containing MEIS2 motifs. OE, overexpression. *p < 0.05, **p < 0.01, ****p < 0.0001 (one-way analysis of variance with Tukey multiple comparisons). Each dot represents one replicate.

Next, since gene expression is regulated by gene regulatory networks (GRNs)^33^ and expression of genes associated with ASD converge in excitatory neurons during these developmental stages in ExN L2-3 IT and ExN L3-5 IT cell subclasses^14, 15^, we inferred cell-type gene regulatory networks (GRNs) for these two cell subclasses using transcriptomic and epigenomic data together with transcription factor (TF) motif binding sites (**Methods**). These cell-type GRNs link regulatory elements with TF binding sites to target genes. We particularly focused on analyzing the subnetworks regulating the genes identified in our developmental maturation trajectory analysis (**Table S4**) and found nine major TFs regulating the driver genes of the two cell subclasses. Among these nine TFs, we identified MEIS2 as a key regulator for ExN L2-3 IT, FOXP2 as a hub TF for ExN L3-5 IT, and MEF2C as an important TF for both ExN subclasses (**Figure 3B**).

Since upper-layer (L2-3) neurons exhibit selective expansion in the primate lineage^23^ and their roles in the cortico-cortical circuits that are important for the unique cognitive and behavioral abilities of primates^30^, we investigated putative regulatory elements that underlie the transcriptomic differences between ExN L2-3 IT and ExN L3-5 IT. We calculated differential accessibility and overrepresented motifs between these neuronal populations throughout midfetal and late-fetal development. Using this strategy, we identified the MEIS2 motif MA0774.1 as one of the overrepresented motifs in the ExN L2-3 IT subclass (**Table S6**). Furthermore, we found that ExN L2-3 IT has both higher *MEIS2* gene expression and a higher MEIS2 motif activity score when compared to ExN L3-5 IT (**Figure 3C**). Because MEIS2 is a transcription factor enriched in upper-layer neurons in primates^10, 23^ and is implicated in neuropsychiatric disorders^32^, we calculated MEIS2 transcription factor footprints across the whole genome in the two cell subclasses and found that, throughout the analyzed developmental periods, ExN L2-3 IT had a higher footprint enrichment than ExN L3-5 IT, with the highest enrichment in ExN L2-3 IT during midfetal development (PCD105-110) and the lowest footprint enrichment in ExN L3-5 IT during late fetal development (PCD155), indicating the selective involvement of MEIS2 in the regulation of target genes in ExN L2-3 IT (**Figure 3C**). Indeed, quantitative immunofluorescence showed that there were significantly fewer MEIS2-expressing cells in L3-5, identified by the expression of *RORB* (**Figure S3B**), compared to L2-3 of dlPFC at PCD155 (**Figure 3D**).

One of the genes potentially regulated by MEIS2 in IT excitatory neurons is the *Nuclear factor I/A* (*NFIA*), which has been shown to be regulated by MEIS2 in retinal progenitor cells^34^ and is linked to callosal hypoplasia, likely reflecting malformations of upper-layer excitatory neurons^35^. By analyzing the distribution of accessible chromatin peaks in the vicinity of *NFIA* across the two neuronal subclasses and developmental periods, we identified 5 active MEIS2 binding sites within 200kb upstream of the *NFIA* transcription start site (**Figure 3E**). We also found three peak-to-gene links (peaks that show higher co-activity with *NFIA* gene expression), with one of the links containing two predicted MEIS2 binding motifs (chr1-163553896-163555161) approximately 181kb upstream of the *NFIA* transcription starting site, suggesting that it may be a potential regulatory element for *NFIA*. To validate this finding, we conducted a dual luciferase assay to assess the activity of the genomic sequence containing the two MEIS2 motifs. Overexpression of MEIS2 resulted in a significant increase in transcription activity driven by the genomic sequence. Furthermore, when we introduced mutations into two potential MEIS2 binding motifs, one of these mutations significantly reduced the activity of the genomic sequence (**Figure 3F**). Taken together, our analyses have unveiled key regulators, such as MEIS2, and gene regulatory networks governing the maturation of distinct IT neuronal populations during midfetal and late-fetal primate brain development.

### Patch-seq of midfetal and late-fetal macaque dlPFC cells identified transcriptomic signatures defining electrophysiological maturation

The midfetal and late-fetal periods are critical stages in cortical development, marked by notable alterations in neuronal morphology, electrophysiology, and gene expression^13^. Although all the analyzed neurons displayed a relatively immature morphology, PCD155 neurons exhibited significantly greater complexity, with greater total dendritic length, increased branching points, and more endings when compared to neurons from PCD100 brains (**Figures S4B-S4F**). To gain a comprehensive understanding of the molecular mechanisms that drive neuronal maturation, we employed Patch-seq, which allows for sequential electrophysiological and transcriptomic analyses of the same cell^36^, in 16 rhesus macaques ranging from PCD85 to PCD155 (**Figures 4A and S4; Table S1**). We used our snRNA-seq data (**Figure 2**) as the reference dataset to infer cell types by mapping the 256 cells profiled by Patch-seq onto the reference UMAP (**Figures 4B and S4I; Table S7**). Approximately 72% of all cells were identified as neurons based on their transcriptomic signature, which included 111 IT neurons (**Table S7**). Our evaluation of the electrophysiological features of three major IT subtypes (L2-3, L3-5, and L6) across different ages revealed that these IT neurons exhibited similar electrophysiological features (**Figures S4J-S4N**) during this developmental window. Therefore, we combined all IT neuron types for downstream analysis.

**Figure 4.**
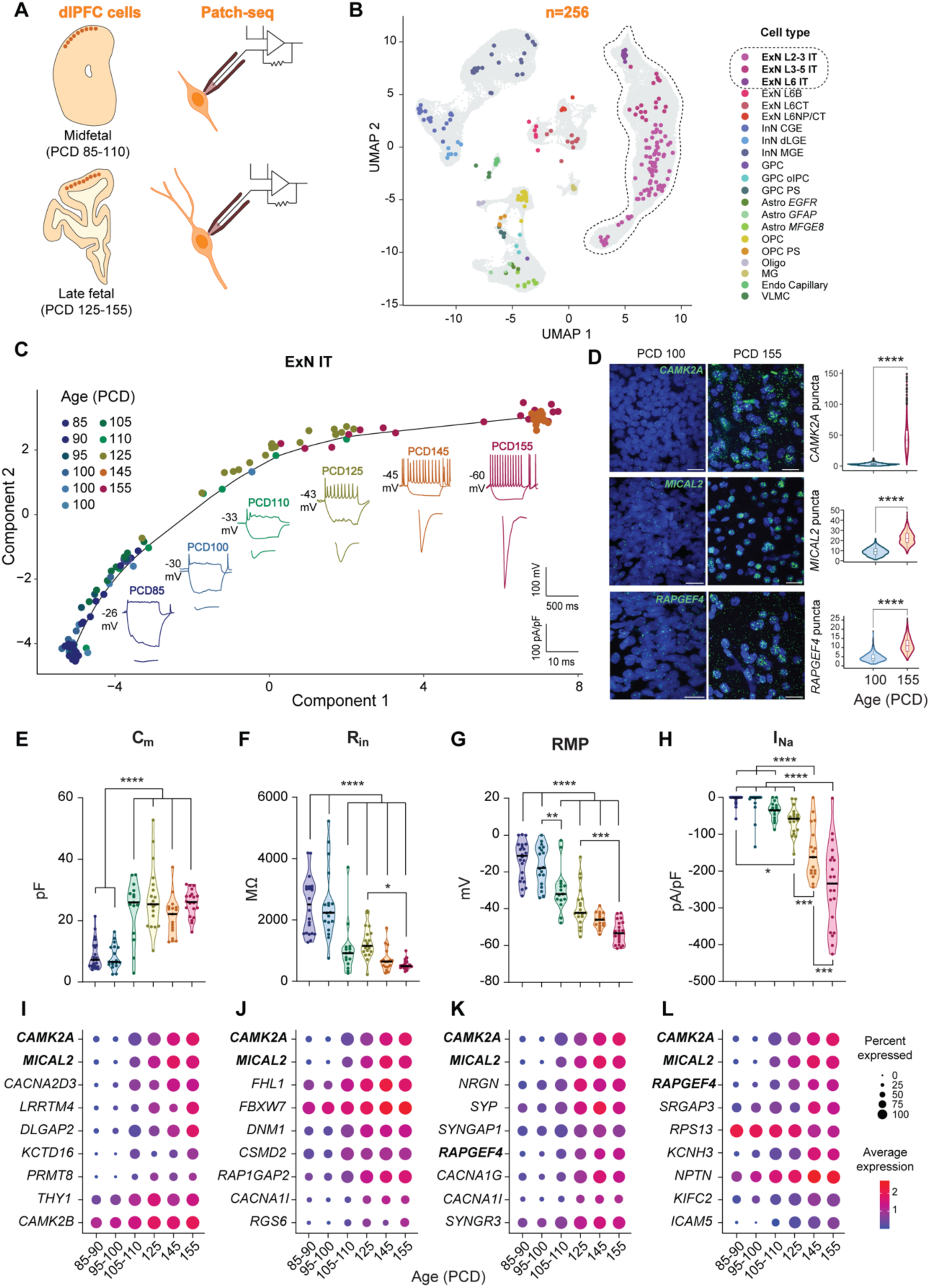
Identification of midfetal and late-fetal neuronal maturation-related genes using Patch-seq. (A) Schematic representation of the Patch-seq experimental design for analyzing midfetal and late-fetal rhesus macaque dlPFC cells. (B) Integration of the 256 cells profiled by Patch-seq onto the snRNA-seq reference UMAP. (C) Pseudotime trajectory based on the gene expression profiles of patched IT neurons from 16 individual rhesus macaque fetal brains at 9 distinct fetal ages, with representative electrophysiological sample traces (voltage responses to depolarizing and hyperpolarizing current injections, and inward sodium currents). (D) In situ hybridization detection and quantification (right panel) of *CAMK2A*, *MICAL2*, and *RAPGEF4* in PCD100 and PCD155 dlPFC. Scale bar, 20 µm. (****p < 0.0001, *t*-test). (E-L) Plots illustrating the developmental changes in electrophysiological properties, including membrane capacitance (C_m_, E), input resistance (R_in_, F), resting membrane potential (RMP, G), and inward sodium current (I_Na_, H) of IT neurons across six different age groups. Bottom panels show the expression levels of C_m_-correlated genes (I), R_in_-correlated genes (J), RMP-correlated genes (K), and I_Na_-correlated genes (L) in IT neurons across the same age groups. (**p < 0.01, ***p < 0.001, ****p < 0.0001, one-way ANOVA, Tukey’s multiple comparisons test).

We employed a pseudotime trajectory analysis to infer a maturation trajectory by integrating differentially expressed genes across age groups in all identified IT neurons (**Methods**). Our analysis revealed a trajectory characterized by the progressive maturation of IT neurons from earlier to later developmental periods (**Figure 4C**). We observed that the proportion of IT neurons exhibiting action potentials (APs) across six age groups (PCD85-90, PCD95-100, PCD105-110, PCD125, PCD145, and PCD155) gradually increased from 0% in the PCD85-90 group to 95% in PCD155, providing strong evidence for electrophysiological maturation (**Figure S4O**). We then assessed neuronal maturation through the lens of the following features that offer unique perspectives on maturation: membrane capacitance (C_m_), input resistance (R_in_), resting membrane potential (RMP), and inward sodium current (I_Na_). Our analysis revealed that, during early midfetal development (from PCD85 to PCD100), none of the electrophysiological features showed significant change (**Figure 4**). We observed a significant increase in C_m_ from PCD100 to PCD105, after which the values remained stable (**Figure 4**). This increase in C_m_ potentially reflects the enlargement of the cell body and more complex cellular structures during the maturation of neurons. In contrast, there was a gradual decline in the averaged value of R_in_ after PCD125 (**Figure 4F**). This effect may be attributed to an increase in the number of ion channels in the neuronal membrane. Moreover, the morphological changes that occur in dendrites (**Figures S4B-S4F**) and spines during maturation may contribute to alterations in signal processing and integration, further enhancing neuronal function. We also observed a conspicuous pattern of increasingly negative RMP from PCD100 to PCD155 (**Figure 4G**), which reflects the developmental maturation in the ion channel composition of these neurons, allowing for improved electrical signal processing and transmission. During neuronal development, the current density of I_Na_ amplifies, thereby increasing the capacity of neurons to generate action potentials. Our results indicate a significant increase in the magnitude of the I_Na_ from PCD105 to PCD155 (**Figure 4H**). This suggests that IT neurons in the dlPFC acquire the ability to initiate and transmit APs after PCD105. In summary, our analyses showed a progression in maturation across four distinct electrophysiological properties. However, it is noteworthy that the temporal dynamics of the four electrophysiological properties were not identical. While RMP and I_Na_ exhibited a progressive increase from PCD95-100 to PCD155, C_m_ and R_in_ displayed drastic changes only at PCD105-110. Our study has uncovered distinct profiles in the temporal dynamics of electrophysiological maturation, where each property exhibits a different rate of maturation.

Similar to IT ExNs, InNs exhibited progressively more negative RMP and I_Na_ from PCD105-110 to PCD155, whereas R_in_ didn’t show any significant differences and C_m_ became higher throughout development (**Figures S5A-S5E**). The proportion of InNs with APs increased from 12.5% in PCD85-90 to 100% in PCD125, PCD145, and PCD155, indicating a faster electrophysiological maturation than ExNs (**Figure S5F**).

We hypothesized that differential expression of distinct genes drives the divergent trends of the four electrophysiological features across maturation. By implementing a correlation analysis between the electrophysiological data and the gene expression levels across all IT neurons, we identified 135, 163, 193, and 142 genes that are associated with the temporal dynamics of electrophysiological features of C_m_, R_in_, RMP, and I_Na_, respectively (**Figure S6A; Table S9**). Importantly, genes that are not expected to exhibit correlation with electrophysiological features, including the housekeeping genes *GAPDH* and *TBP*, showed no significant correlation with any of the electrophysiological features (**Figure S6B**). Several genes, including *CAMK2A* and *MICAL2*, exhibited high correlation with all four investigated electrophysiological features (**Figures 4I-4L**). Other genes, such as *RAPGEF4*, exhibited significant correlation with one or two of the electrophysiological features. To validate our Patch-seq data, we performed in situ hybridization on dlPFC sections of PCD100 and PCD155 macaques for select genes that demonstrated significant correlation with all (*CAMK2A*, *MICAL2*) or some (*RAPGEF4*, *NRGN*) of the electrophysiological features. Consistent with the Patch-seq results, the expression levels of all tested genes were significantly higher in PCD155 compared to PCD100 (**Figures 4D and S6C**).

Therefore, our Patch-seq analysis of dlPFC neurons across midfetal and late-fetal development revealed evidence of neuronal maturation across three distinct aspects: morphological, transcriptomic, and electrophysiological properties, and identified genes associated with maturation of specific electrophysiological features.

### *RAPGEF4* is crucial for the morphological and electrophysiological maturation of excitatory neurons

Our Patch-seq analysis demonstrated a marked upregulation of *RAPGEF4* expression in IT neurons during cortical development (**Figures 4K-4L**), which was also corroborated by our snRNA-seq data (**Figure 5A**). RAPGEF4 (Rap Guanine Nucleotide Exchange Factor 4), also known as EPAC2 (exchange protein directly activated by cyclic AMP 2), is a regulator of small GTPase signaling and a protein kinase A (PKA)-independent cAMP target^37–39^. We found that *RAPGEF4* expression exhibited a high degree of correlation with various electrophysiological features, with the highest correlation observed with I_Na_ (r = −0.57) and the second highest correlation with RMP (r = −0.47) (**Figure S6E**). To confirm that RAPGEF4 is required for neuronal maturation, we infected ex vivo macaque brain slices with lentivirus (LV) expressing mScarlet and shRNA against *RAPGEF4* (LV-sh*RAPGEF4*) together with LV expressing mNeonGreen and control shRNA (LV-sh*NC*) (**Figure 5B**). The neurons with *RAPGEF4* knockdown (sh*RAPGEF4*, mScarlet^+^) exhibited reduced dendritic complexity, total dendritic length, branching points, and number of endings compared to controls (mNeonGreen^+^mScarlet^-^) (**Figures 5C-5H**). In addition, LV-sh*RAPGEF4*-infected neurons exhibited smaller I_Na_ (**Figures 5I and 5M**) and less negative RMP (**Figures 5I and 5L**), without significant changes in C_m_ and R_in_, compared to control neurons (**Figures 5J-5K; Table S10**). These results indicate that a deficiency of *RAPGEF4* in neurons leads to delayed maturation of I_Na_ and RMP, which is consistent with our finding that *RAPGEF4* had a strong correlation with I_Na_ and RMP (**Figures 4K-4L and S6E**).

**Figure 5.**
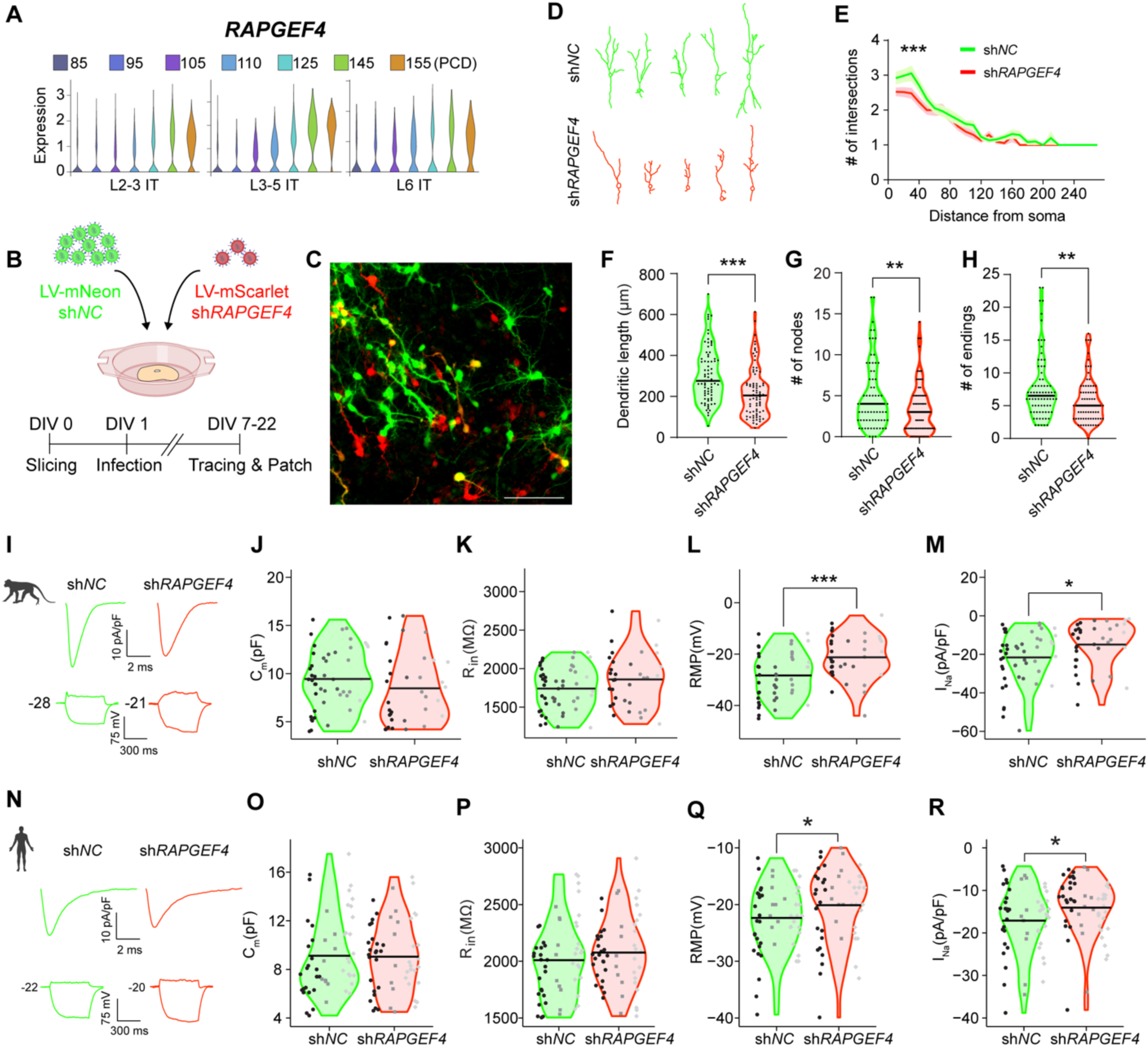
Knockdown of *RAPGEF4* impairs morphological and electrophysiological maturation of cortical neurons in macaque organotypic dlPFC slices. (A) Violin plots showing the expression of *RAPGEF4* in IT neurons from PCD85 to PCD155. (B) Schematic of the experimental setup for investigating the morphological and electrophysiological alterations after *RAPGEF4* knockdown in ex vivo dlPFC slices. (C) Representative image depicting *RAPGEF4* knockdown (both mScarlet^+^[red] and mScarlet^+^/mNeon^+^[yellow]) and control (mNeon^+^[green only]) neurons in macaque dlPFC slices. Scale bar, 100 μm. (D) Sample traces depicting the morphology of both *RAPGEF4* knockdown (bottom panel) and control (top panel) neurons. (E) Sholl analysis of dendritic complexity in *RAPGEF4* knockdown neurons compared to control neurons [F(1,150) = 14.531, ***p < 0.001, multivariate analysis of variance]. (F-H) Knockdown of *RAPGEF4* decreases total dendritic length (F), number of nodes (G), and number of endings (H) (*t*-test, ***p < 0.001, **p < 0.01). (I) Representative traces of inward sodium currents and voltage responses to depolarizing and hyperpolarizing current injections for macaque *RAPGEF4* knockdown (right panel) and control (left panel) neurons. (J-M) Knockdown of *RAPGEF4* in macaque organotypic slices leads to reduced I_Na_ (M) and less negative RMP (L), without significantly change in C_m_ (J) and R_in_ (K) (*t*-test, ***p < 0.001, *p < 0.05). Each dot represents one patched cell. All cells were collected from four individual fetal brains, with each fetal brain indicated by a different color: black, dark gray, medium gray, and light gray. (N) Representative traces of inward sodium currents and voltage responses to depolarizing and hyperpolarizing current injections for human *RAPGEF4* knockdown (right panel) and control (left panel) neurons. (O-R) Knockdown of *RAPGEF4* in human organotypic slices leads to reduced I_Na_ (R) and less negative RMP (Q), without significantly change in C_m_ (O) and R_in_ (P) (*p < 0.05). Each dot represents one patched cell. All cells were collected from four individual fetal brains, with each fetal brain indicated by a different color: black, dark gray, medium gray, and light gray.

RAPGEF4 is enriched in neurons compared to other cell types^39^. The levels of *RAPGEF4* increase in the human cortex throughout development, compared to other brain regions or organs^9, 40^. Based on our observations in macaque, we hypothesized that *RAPGEF4* expression is also associated with the maturation of electrophysiological properties in human. We infected human midfetal cortical brain slices with LV-sh*RAPGEF4* and performed patch-clamp recording. As in macaque, we observed reduced I_Na_ and RMP, but not C_m_ and R_in_, in LV-sh*RAPGEF4*-infected compared to LV-sh*NC*-infected neurons (**Figures 5N-5R; Table S10**). To confirm that the observed effect was not caused by off-target effects of the shRNA, we replicated the experiment and observed the same results using a different shRNA targeting *RAPGEF4* in human neurons (**Figures S6H-S6L**). The role of RAPGEF4 in regulating RMP and I_Na_ maturation in both macaque and human neurons is consistent with the functional enrichment for “regulation of GTPase activity” in genes driving neuronal maturation (**Figure S6D**). These data validate our approach to identify genes that drive the maturation of specific electrophysiological features and reveal the novel role of *RAPGEF4* in morphological and electrophysiological maturation of neurons in both macaque and human cortical development.

### *CHD8* knockdown leads to impaired maturation of human cortical excitatory neurons

Recent advances in genomics have revealed important roles of chromatin regulators during midfetal PFC development and implicated them in neurodevelopmental disorders^41, 42^. CHD8 (Chromodomain Helicase DNA-binding protein 8) is a chromatin remodeling protein that plays a crucial role in regulating gene expression during brain development and has been linked to ASD^42–44^. To investigate the role of *CHD8* during human midfetal cortical development, a critical developmental period for ASD^14, 15^, we knocked down *CHD8* expression in ex vivo human midfetal brain slices using LV expressing sh*CHD8* and mScarlet together with LV expressing *shNC* and mNeonGreen (**Figures 6A and S7B-S7C**). LV-sh*CHD8*-infected neurons displayed significantly shorter dendrites compared to control neurons (**Figures 6B, 6D-6E and S7D-S7F**), suggesting that a deficiency in *CHD8* disrupts morphological maturation of cortical neurons during midfetal development, which is consistent with previous findings in mice^44^.

**Figure 6.**
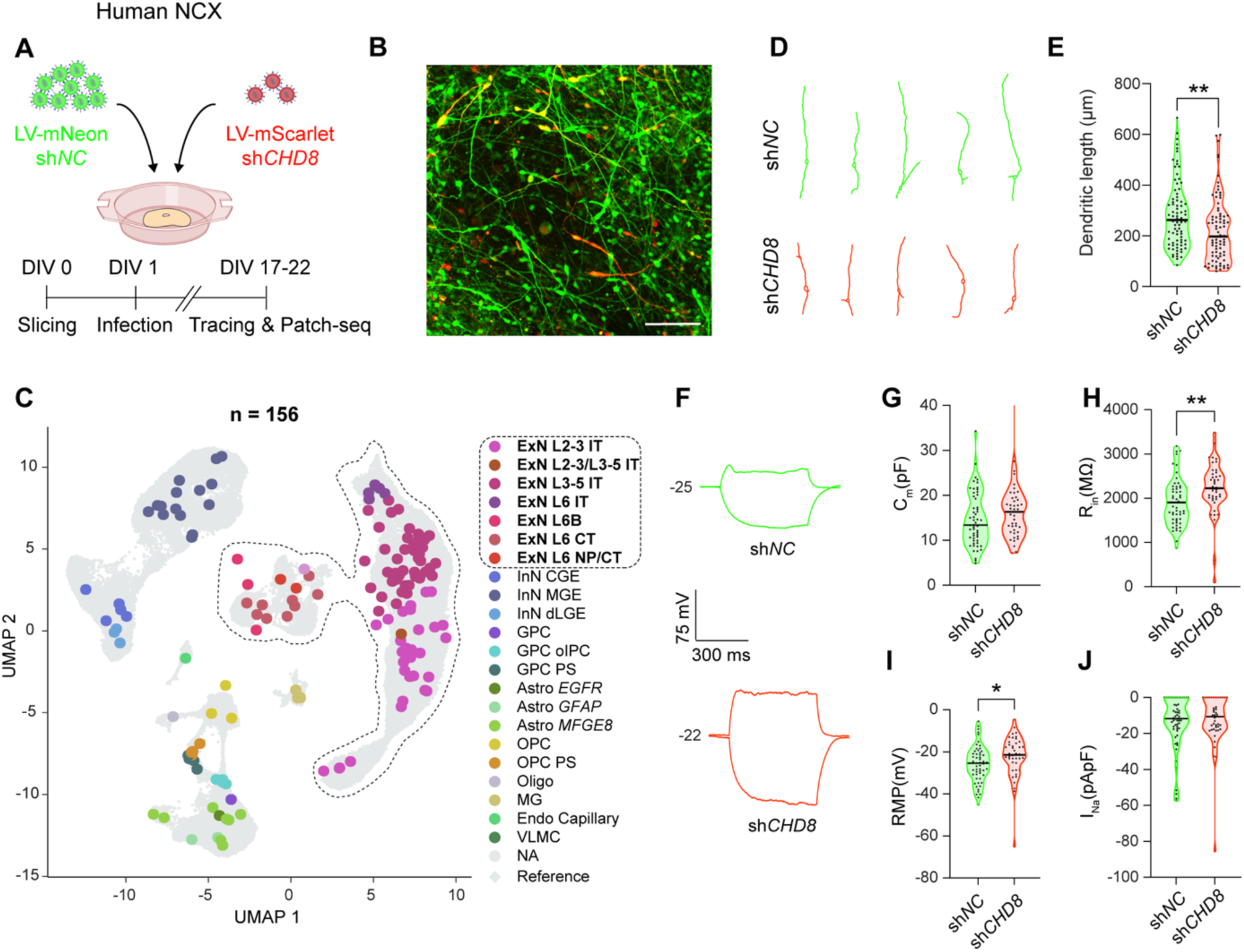
Knockdown of *CHD8* impairs maturation of cortical excitatory neurons. (A) Schematic diagram illustrating the experimental setup used to study the morphological and electrophysiological changes after knockdown of *CHD8* in ex vivo human fetal cortical slices. (B) Representative image showing *CHD8* knockdown (mScarlet^+^[red], mScarlet^+^ and mNeon^+^[yellow]) and control (mNeon^+^[green]) neurons in human cortical slices. Scale bar, 100 μm. (C) Integration of the 156 cells profiled by Patch-seq onto the snRNA-seq reference UMAP. (D) Sample traces showing the morphology of both *CHD8* knockdown and control neurons. (E) *CHD8* knockdown reduces total dendritic length (**p < 0.01, *t*-test). (F) Representative traces (voltage response to depolarizing [50 pA] and hyperpolarizing [-50 pA] current injections) for control neurons and *CHD8* knockdown. (G-J) *CHD8* knockdown increases R_in_ (H) and RMP (I) but does not significantly alter C_m_ (G) and I_Na_ (J) (*p < 0.05, **p < 0.01, *t*-test).

We then performed Patch-seq on 156 cells from slices that contained both *CHD8*-knockdown and control cells (**Figures 6A and S7G**). To determine the cellular identity of recorded cells, we compared their transcriptome with the reference snRNA-seq data and assigned each queried cell a specific cell type (**Figures 6C and S7H; Table S11**). Out of the 156 cells examined, there were 55 excitatory neurons infected with LV-sh*NC* and 42 infected with LV-sh*CHD8* (**Table S11**). LV-sh*CHD8*-infected excitatory neurons had less negative RMP and a significant increase in R_in_ (**Figures 6I and 6H**) but not in C_m_ and I_Na_ (**Figures 6G, 6J and S7I**), compared to the controls. To rule out the possibility that the observed effect is due to shRNA off-target effects, we repeated the experiment with an alternative shRNA in human cultured neurons and observed similar deficits in both R_in_ and RMP (**Figures S7J-S7N**). Therefore, *CHD8* knockdown produces a discernible impact on the electrophysiological properties of midfetal human neurons, which is consistent with its role during early development^43, 44^.

### RAPGEF4 rescues electrophysiological deficits of CHD8-deficient human cortical neurons

To determine which genes might underlie the observed differences in R_in_ and RMP maturation caused by *CHD8* knockdown, we identified the differentially expressed genes between LV-sh*CHD8* and LV-sh*NC* and intersected this list with the genes that were associated with the maturation of these two electrophysiological features (**Table S9**). We found 10 genes associated with R_in_, 11 genes associated with RMP, and 20 genes associated with both (**Figure 7A**). These included genes that are associated with the overall maturation of neurons, such as *NRGN* and *CACNA1I*, the R_in_-associated *RGS6*, a GTPase regulator important for activity-dependent morphological and electrical maturation^45^, and *RAPGEF4*, which we found to significantly impact the maturation of RMP (**Figures 7B-7E and 5**).

**Figure 7.**
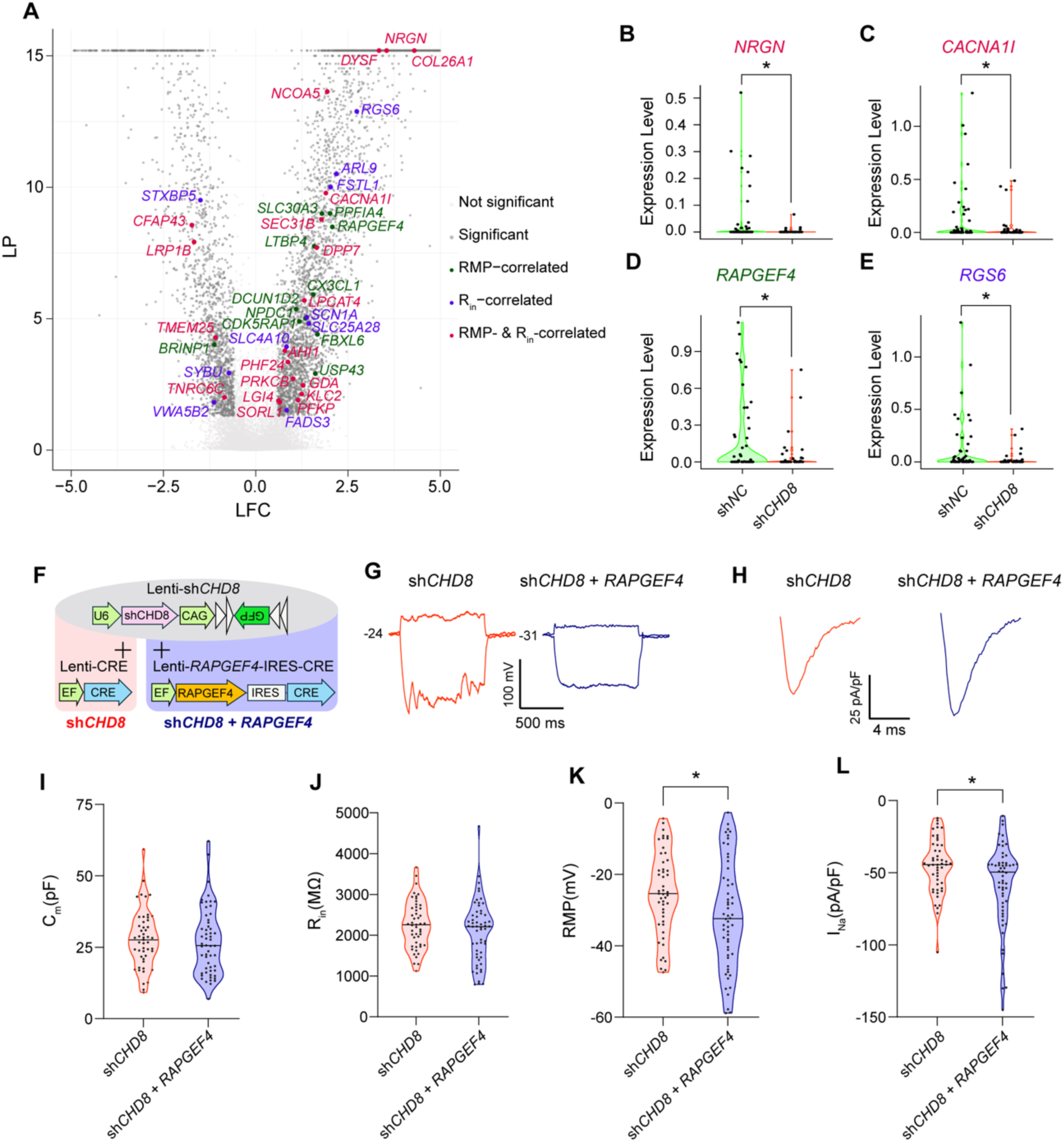
Exogenously expressed RAPGEF rescues neuronal maturation deficits resulting from CHD8 deficiency. (A) Volcano plot depiction (-log_10_(*P_adj)_*) vs log_2_(*fold-change*)) of differentially expressed genes between LV-sh*NC* and LV-sh*CHD8* -infected excitatory neurons. Genes with expression profiles that significantly correlate with the RMP (green), R_in_ (purple), or both (red) electrophysiology features are labeled. (B-E) Violin plots showing the expression levels of *NRGN* (B), *CACNA1I* (C), *RAPGEF4* (D), and *RGS6* (E), highlighting the effect of *CHD8* knockdown on these neuronal maturation genes (Figures 4I-4L). (F) Schematic of lentiviral vectors used for knocking down *CHD8* and overexpression of *RAPGEF4* in human cortical ex vivo organotypic slices. (G) Representative traces (voltage response to depolarizing [50 pA] and hyperpolarizing [-50 pA] current injections) for neurons with *CHD8* knockdown and neurons with *CHD8* knockdown and *RAPGEF4* overexpression. (H) Representative traces of inward sodium currents to depolarizing current injections for neurons with *CHD8* knockdown and neurons with *CHD8* knockdown and *RAPGEF4* overexpression. (I-L) *RAPGEF4* rescues RMP (K) and I_Na_ (L) but does not significantly alter C_m_ (I) and R_in_ (J) of neurons with *CHD8* knockdown (*p < 0.05, **p < 0.01, *t*-test).

To determine whether reduced RAPGEF4 levels might contribute to the observed reduced RMP in *CHD8* deficient neurons, we expressed *RAPGEF4* in human cortical neurons that were knockdown for *CHD8* (sh*CHD8*) (**Figure 7F**). Exogenously expressed *RAPGEF4* restored normal maturation of both RMP and I_Na_, with no significant effect on R_in_ or C_m_, in *CHD8*-deficient neurons (**Figures 7G-7L; ; Table S12**). Therefore, *CHD8* deficiency in primate cortical neurons leads to reduced expression of genes important for electrophysiological maturation, including *RAPGEF4*, which may contribute to neuronal maturation deficits.

## DISCUSSION

In this study, we employed multimodal approaches including snMultiome and Patch-seq to interrogate mechanisms driving neuronal maturation in the primate dlPFC during midfetal and late-fetal development. We identified dynamic gene expression changes across cell types in their maturation trajectories as well as critical developmental periods for the maturation of specific electrophysiological properties in excitatory and inhibitory neurons.

We confirmed that, based on their molecular profiles, excitatory neurons gain their laminar and projection identities earlier than interneuronal final specification. This corroborates previous findings that pointed to the relevance of extrinsic factors to the final specification of interneurons during perinatal development^20, 45^. We focused our analyses on the IT excitatory neurons due to their relevance both to primate brain evolution and their association to neuropsychiatric disorders. Our integrative analysis of gene expression and chromatin accessibility data unveiled gene regulatory networks associated with neuron maturation of both excitatory and inhibitory neurons and highlighted MEIS2 as a key transcriptional factor for L2-3 IT neuronal development and maturation.

By integrating electrophysiology, morphology, and functional genomics analyses of primate dlPFC development, we discovered that different electrophysiological properties mature at different rates. While RMP change gradually and linearly during maturation, C_m_ and R_in_ exhibit a jump at the end of neurogenesis during midfetal developmental, I_Na_ exhibited significant maturation only at the late-fetal period, which coincides with a significant increase in the number of neurons able to fire action potentials. The distinct rate of maturation among electrophysiological properties reveals the intricate and finely tuned regulation of neuronal maturation during midfetal and late-fetal development of primate PFC.

Our Patch-seq approach allowed us to investigate the genes associated with the maturation of specific electrophysiological features. We have uncovered genes, including kinases, small GTP signaling molecules, neurotransmitter receptors, ion channels, and transporters that are associated with maturation of specific electrophysiological features that likely underlie the differences we observed in their maturation kinetics. One of these genes is *RAPGEF4*, which we have identified as being significantly correlated with the maturation of resting membrane potentials and inward sodium currents. RAPGEF4, a guanine exchange factor in small GTPase pathway, is a direct target of cAMP and has been shown to regulate dendritic spine remodeling in mice and its deficiency in mice leads to learning and memory deficits^39, 46^. However, its role in primate brain development or human disease is unknown. Dysregulation of the cAMP pathway has been implicated in a wide range of neurological disorders and a potential treatment to elevate cAMP levels is currently in clinical trial for fragile X syndrome^47–49^. The small GTPase pathway has been shown to have critical roles during maturation of human neurons^50^. Our study demonstrates that RAPGEF4 may be one of the small GTPase regulators driving neuronal maturation during primate brain development, especially for RMP, an electrophysiological feature regulated by GTPase signaling.

Finally, to demonstrate the potential of our experimental platform in exploring the developmental origin of human diseases, we showed that knocking down of a high-confidence ASD gene – *CHD8* – impaired morphological and electrophysiological maturation of excitatory neurons in human organotypic slices. This impairment was driven by the downregulation of the expression of several key genes that we identified to be relevant for the electrophysiological maturation of neurons, including *RAPGEF4 and RGS6*, two regulators of small GTPase signaling critical for neuronal maturation^50^. By restoring the expression of *RAPGEF4* in CHD8-deficient neurons we were able to rescue the deficits in electrophysiological maturation of these neurons. The chromatin regulator CHD8 is expressed at high levels during early-fetal and midfetal development and the levels of CHD8 do not exhibit significant upregulation during midfetal to late-fetal neuronal maturation. How CHD8 and other early expressing chromatin regulators converging at midfetal development contribute to ASD remains unclear. Our finding sheds light on how a chromatin remodeler may regulate the physiological maturation of neurons, thus impairing their normal functioning.

Together, our study has revealed key genes and gene networks driving neuronal maturation during a critical period of prefrontal development in primates and linked the function of a novel regulator of the maturation of specific electrophysiological features with a high-confidence ASD risk gene. Our results demonstrate the power of our multimodal approach and its application for future studies aiming at understanding how genes regulate neuronal maturation and its implication in brain disorders. Finally, we summarized our results including cell clusters, genes and networks in an open-access, searchable web app, which will allow researchers to investigate the role of different genes on neuronal maturation.

## ACKNOWLEDGEMENTS

We thank Y. Xing, S. Krebsbach, D. Phan, M. Weidenfeller, and H. Thurston for technical assistance; K. Knobel at the Waisman IDD Model Core for core services; Dr. Heather Simmons, Director of Pathology Services (WNPRC) for non-human primate tissue acquisition; and the Birth Defects Research Laboratory (BDRL) at the University of Washington for human tissue acquisition. This work was supported by the National Institutes of Health (R01MH118827 and R01NS105200 to X.Z.; R01HD064743 to Q.C; R01NS064025, R01AG067025, and RF1MH128695 to D.W; 1R01HD106197 and UM1MH130991 to A.M.M.S; P51 OD011106 to WNPRC; P50HD105353 to Waisman Center; and R24HD000836 to BDRL). Further support was provided by the DOD IIRA grant (X.Z.), SFARI pilot grant (X.Z., A.M.M.S., Q.C., and D.W.), Jenni and Kyle Professorship (X.Z.), Brain and Behavior Research Foundation (29721 to A.M.M.S.), Brain Research Foundation (BRFSG-2023-11 to A.M.M.S.), and the Medical Scientist Training Program T32 (GM140935 to R.D.R.).

## AUTHOR CONTRIBUTIONS

Y.G., Q.D., Q.C., X.Z., and A.M.M.S. conceived and designed the study. Y.G., Q.D., R.D.R., M.S., D.K.S., and A.M.M.S. performed the experiments. K.H.A., J.S., T.J., S.L., S.K., and D.W. analyzed the genomics data. J.E.L., D.D., I.G. provided the analyzed specimens. Y.G., Q.D., K.H.A., Q.C., X.Z., and A.M.M.S. wrote the manuscript. All authors edited the manuscript.

## RESOURCE AVAILABILITY

The genomic data has been submitted to GEO (GSE235493). The data can be interactively visualized at https://daifengwanglab.shinyapps.io/DevMacaquePFC/. The code and data used for generating figures can be accessed at Zenodo: https://zenodo.org/records/11123264. All other data are available in the main article or supplemental information. Further information and requests for resources should be directed to the corresponding authors.

